# Identifiability of Tissue Material Parameters from Uniaxial Tests using Multi-start Optimization

**DOI:** 10.1101/2020.09.09.289702

**Authors:** Babak N. Safa, Michael H. Santare, C. Ross Ethier, Dawn M. Elliott

## Abstract

Determining tissue biomechanical material properties from mechanical test data is frequently required in a variety of applications, e.g. tissue engineering. However, the validity of the resulting constitutive model parameters is the subject of debate in the field. Common methods to perform fitting, such as nonlinear least-squares, are known to be subject to several limitations, most notably the uniqueness of the fitting results. Parameter optimization in tissue mechanics often comes down to the “identifiability” or “uniqueness” of constitutive model parameters; however, despite advances in formulating complex constitutive relations and many classic and creative curve-fitting approaches, there is no accessible framework to study the identifiability of tissue material parameters. Our objective was to assess the identifiability of material parameters for established constitutive models of fiber-reinforced soft tissues, biomaterials, and tissue-engineered constructs. To do so, we generated synthetic experimental data by simulating uniaxial tension and compression tests, commonly used in biomechanics. We considered tendon and sclera as example tissues, using constitutive models that describe these fiber-reinforced tissues. We demonstrated that not all of the model parameters of these constitutive models were identifiable from uniaxial mechanical tests, despite achieving virtually identical fits to the stress-stretch response. We further show that when the lateral strain was considered as an additional fitting criterion, more parameters are identifiable, but some remain unidentified. This work provides a practical approach for addressing parameter identifiability in tissue mechanics.

**Statement of Significance:** Data fitting is a powerful technique commonly used to extract tissue material parameters from experimental data, and which thus has applications in tissue biomechanics and engineering. However, the problem of “uniqueness” or “identifiability” of the fit parameters is a significant issue, limiting the fit results’ validity. Here we provide a novel method to evaluate data fitting and assess the uniqueness of results in the tissue mechanics constitutive models. Our results indicate that the uniaxial stress-stretch experimental data are not adequate to identify all the tissue material parameters. This study is of potential interest to a wide range of readers because of its application for the characterization of other engineering materials, while addressing the problem of uniqueness of the fitted results.

## 1 Introduction

Knowledge of material properties is valuable to study complex physio-mechanical tissue function, to monitor pathophysiological changes, and to characterize tissue-engineered constructs [1–5]. Despite the widespread use of parameter optimization when fitting experimental data, there are important limitations to this technique arising from the uniqueness (or lack thereof) of the fitted parameters. Therefore, a systematic and practical understanding of this limitation in parameter optimization is needed.

Several optimization approaches have been used in tissue mechanics, yet their success is a subject of debate. Nonlinear least-squares optimization (NLSQ) is perhaps the most commonly used approach. However, NLSQ methods are prone to local minima traps, and depend strongly on the initial guesses used in the fitting algorithm; further, when fitting parameters of highly complex, nonlinear, and anisotropic tissues, uniqueness (or identifiability) is another hurdle [6]. These problems limit the value of the resulting fitted parameters. Global optimization algorithms, such as genetic algorithms, particle swarm, and simulated annealing, have been used in tissue mechanics to avoid local minima trap with variable success [5,7,8].

Parameter optimization in tissue mechanics is often a problem in the “identifiability” or “uniqueness” of constitutive model parameters [9,10]. A general definition for identifiability is provided in the classic text by Walter and Pronzato [10]. Several studies have previously addressed identifiability of tissue material parameters. For example, Hartmann and Gilbert used the determinant of the Hessian matrix to study identifiability of the two parameters (bulk and shear modulus) describing an elastic material using analytical and finite element solutions [11]; in another study, Akintunde and co-workers used rank deficiency of the Fisher information matrix to study uncertainty and identifiability in murine patellar tendon stress-strain responses [12]. Despite advances in formulating complex constitutive relations and curve-fitting approaches, there is currently no accessible framework to assess identifiability of tissue material parameters.

The objective of this study was to assess the identifiability of material parameters for established constitutive models used to describe the anisotropic and nonlinear response of fiber-reinforced soft tissues, biomaterials, and tissue engineered constructs. We used a numerical approach and focused on several commonly-performed canonical experiments: uniaxial tension and unconfined compression. We used a Monte-Carlo-type multi-start optimization approach [13,14] that Safa and co-workers recently implemented to study poroelasticity and inelasticity of tendon [15,16]. This method enables exploration of the search space around the initial guess and reveals parameter sets that produce the same mechanical response. We show that while some parameters are identifiable, many are not, even though nearly perfect fits to the stress-stretch response can be achieved. We further show that when we impose a second fitting criteria, namely lateral strain, more parameters are identifiable, but some are still not.

## 2 Methods

### 2.1 Overview

An overview of the methods is shown in Figure 1. We first specified constitutive models for nonlinear isotropic and fiber-reinforced anisotropic materials (Section 2.2, Table 1). We numerically implemented these models using the kinematics of uniaxial tension and compression with traction-free lateral boundary conditions (Section 2.3). We next used representative baseline material parameters from sclera in compression and tendon in tension to simulate the baseline “experimental” tissue stress-stretch data for each constitutive model (Section 2.4, Table 1). We then used a multi-start least-squares optimization curve-fit to the baseline response with a wide search space and 600 random initial guesses per material parameter, and assessed the quality of the fits compared to the baseline stress-stretch response (Section 2.5). Because we observed that several parameter sets could reproduce the stress-stretch response, we added a second assessment criteria based on the quality of the lateral strain prediction (Section 2.6). Finally, we assessed the identifiability of each material parameter for each constitutive model by comparing the fitted parameter values to the baseline parameter values (Section 2.7).

**Table 1:**
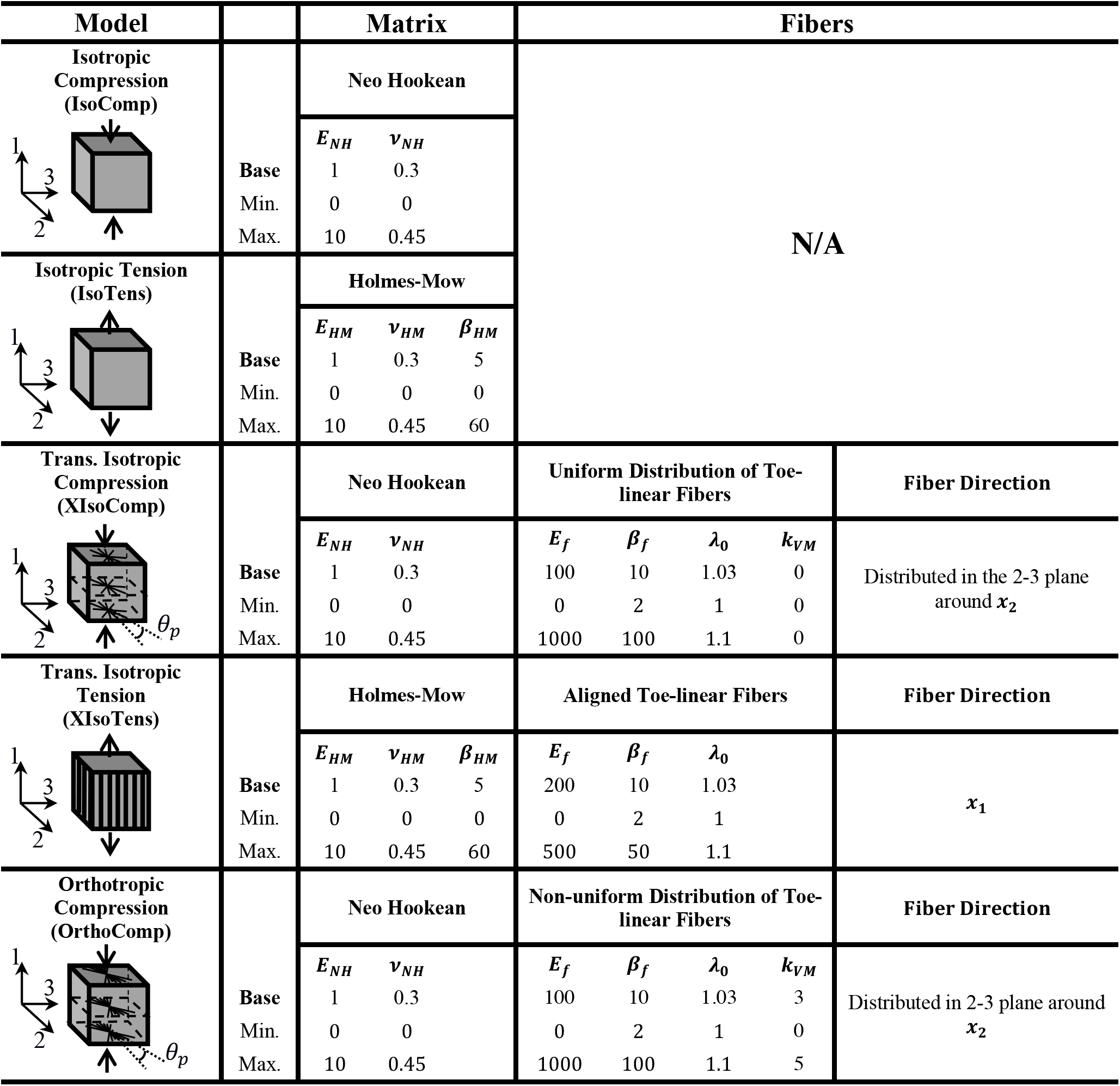
Summary of the model parameters, including baseline parameter values and ranges. In each model, the moduli are non-dimensional

**Figure 1:**
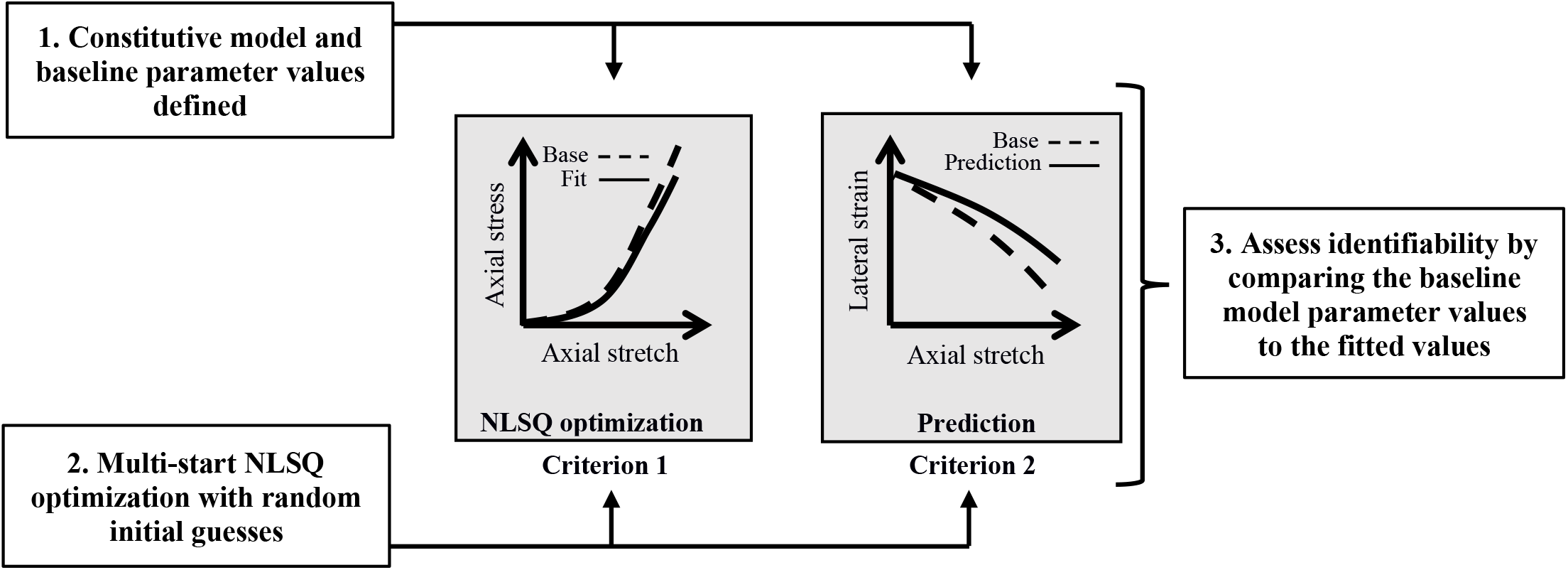
Schematic overview of the methods to assess the identifiability of model parameters.

### 2.2 Constitutive Models

Constitutive models for nonlinear isotropic and fiber-reinforced anisotropic materials that are loaded in uniaxial tension or compression were taken from well-established models [17–22]. We specifically considered two isotropic material models: neo-Hookean in compression and Holmes-Mow in tension, based on typical usage. We added fibers to these isotropic materials to achieve transverse isotropy or orthotropy. The constitutive models, derived from the Helmholtz free energy, are provided in Appendix A. The material parameters are briefly described below.

#### 2.2.1 Isotropic models

For the isotropic model in compression (IsoComp) we used a compressible neo-Hookean material description [21], with the subscript “NH” being used throughout to refer to the neo-Hookean constitutive material parameters. The neo-Hookean constitutive relation has two material parameters (Table 1): *E*_*NH*_ is the Young’s modulus, and *v*_*NH*_ is Poisson’s ratio.

For the isotropic model in tension (IsoTens) we used a Holmes-Mow material description [22], with the subscript ‘HM’ being used throughout to refer to the Holmes-Mow constitutive material parameters. This constitutive model has three material parameters (Table 1): *E*_*HM*_ is the Young’s modulus, *v*_*HM*_ is Poisson’s ratio, and *β*_*HM*_ is the nonlinearity parameter (*β*_*HM*_ > 0).

#### 2.2.2 Transversely isotropic models

For the transversely isotropic model in compression (XIsoComp), we embedded a continuous fiber distribution [20] in a compressible neo-Hookean matrix, More specifically, the fibers were uniformly distributed in the plane perpendicular to the applied loading direction. The fiber constitutive relation was defined by the well-established toe-linear stress-stretch response [17–19], where the stress is nonlinear in the toe-region until reaching the transition stretch, *λ*_0_, after which it is linear with a fiber modulus *E*_*f*_, where the subscript ‘*f*’ stands for fibers. There were five material parameters in the XIsoComp model (Table 1): *E*_*NH*_, *v*_*NH*_ for the matrix (section 2.2.1) and *E*_*f*_, *β*_*f*_, and *λ*_0_ for the in-plane fibers, where *β*_*f*_ (*β*_*f*_ ≥ 2) is the fiber nonlinearity parameter.

For the transversely isotropic model in tension (XIsoTens), we embedded fibers in a Holmes-Mow matrix. The fibers had the same toe-linear response as in XIsoComp, and they were oriented parallel to the loading direction. As a result, there were six material parameters in the XIsoTens model: *E*_*HM*_, *v*_*HM*_, *β*_*HM*_ for the matrix (section 2.2.1) and *E*_*f*_, *β*_*f*_, *λ*_0_ for the fibers (Table 1).

#### 2.2.3 Orthotropic model

For the orthotropic model in compression (OrthoComp) we used the same formulation as the XIsoComp model, except that the in-plane fiber distribution was assumed to be nonuniform, with the modal fiber orientation set to be along axis 2 (i.e., *θ*_*p*_ = 0). This resulted in an orthotropic symmetry in the material configuration, which is characteristic of peripapillary sclera [23,24]. The OrthoComp model has seven material parameters: *E*_*NH*_, *v*_*NH*_ for the matrix and *E*_*f*_, *β*_*f*_, *λ*_0_ and *k*_*VM*_ for the fibers, where *k*_*VM*_ > 0 characterizes the in-plane alignment of fibers.

### 2.3 Kinematics and boundary conditions

The constitutive models were numerically implemented using the kinematics of uniaxial compression or tension with traction-free lateral boundary conditions. We chose these kinematic conditions since they represent canonical experiments frequently used to determine material properties. Uniaxial deformation was described by specifying the deformation gradient tensor (**F**) to be

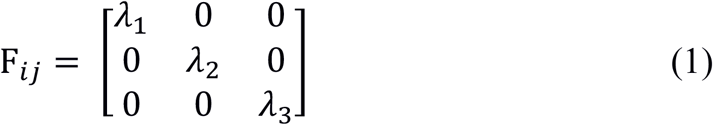

where the {*λ*_*i*_} are the stretches in the directions of the basis vectors (***e***_***i***_; *i* = 1 … 3). Off-diagonal elements of **F** in uniaxial deformation and compression are zero, therefore the reference configuration corresponded to *λ*_*i*_ = 1 (zero strain). To model tension, *λ*_1_ was allowed to vary over the interval [1,1.2], while to model compression, the interval was *λ*_1_ ∈ [0.8, 1]. The values for *λ*_2_ and *λ*_3_ were calculated by imposing traction-free lateral boundary conditions, i.e.,

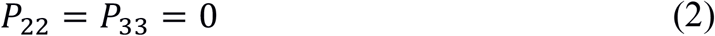

where *P*_22_ and *P*_33_ are components of the first Piola-Kirchhoff stress on lateral surfaces (Figure 2).

**Figure 2:**
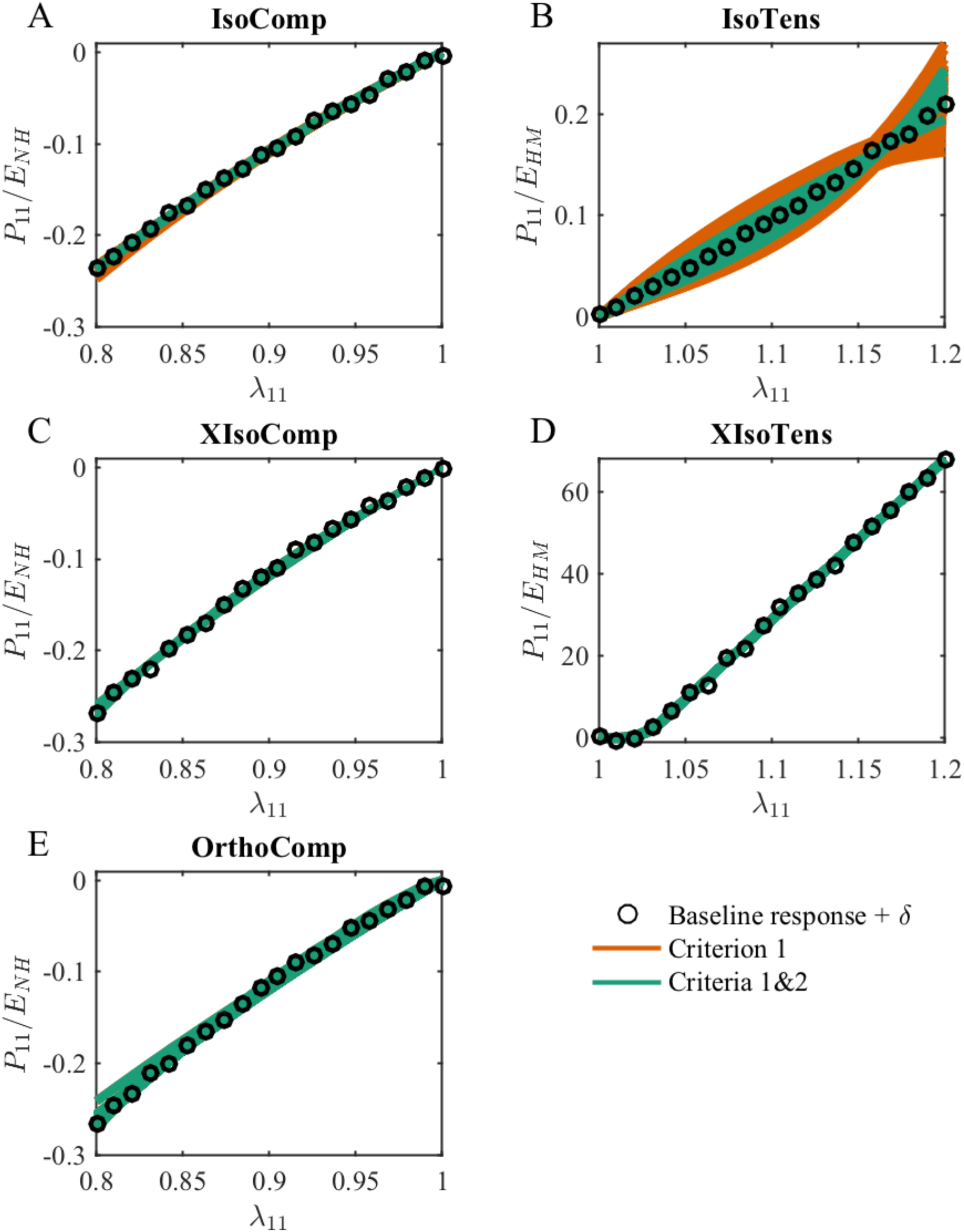
The accepted fits arising from different initial guesses based on only Criterion 1 (orange lines) and Criteria 1&2 (green lines) for the (A) IsoComp, (B) IsoTens, (C) XIsoComp, (D)XIsoTens, and (E) OrthoComp models. All the model fits closely matched their corresponding baseline (“experimental”) stress response, and in most cases, the fits essentially fully overlapped. However, for each model, the fitted stress responses were generated with different sets of parameters. The plotted quantity is the axial component of the Piola-Kirchhoff stress normalized by the baseline parameter value of the matrix modulus. The symbols show the baseline stress with added noise, *δ*,

In order to calculate the stress-stretch response (for both the baseline response and during optimization), we discretized *λ*_1_ into 20 steps over the specified ranges, and for each step we incremented (for tension) or decremented (for compression) *λ*_1_ starting at *λ*_1_ = 1. At each value of *λ*_1_, a nonlinear least squares optimization method was used to calculate values of 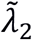 and 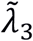 that force the “22” and “33” components of the trial stress tensor 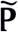 to zero by requiring

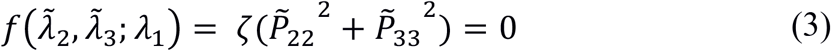

where *ζ* was a penalty factor (*ζ* = 10^6^) to improve stability of the optimization stability and had no effect on the final values of *λ*_2_ and *λ*_3_. Note that this use of optimization to calculate *λ*_2_ and *λ*_3_ is separate from the parameter optimization. Finally, we used the standard continuum mechanics formula (**P** = *∂*Ψ/*∂***F**) to calculate the Piola-Kirchhoff stress from the Helmholtz free energy expressions (Appendix A).

### 2.4 Baseline parameter values

Representative material parameters from sclera in compression and tendon in tension were used to create the baseline “experimental” tissue stress-stretch data for each constitutive model (Table 1). Specifically, the baseline parameter values were chosen to represent peripapillary sclera for the compression models (IsoComp, XIsoComp, and OrthoComp), and tendon for the tension models (IsoTens, and XIsoTens). The only dimensional parameters in the constitutive relations were the Young moduli (*E*_*NH*_, *E*_*HM*_, and *E*_*f*_). We nondimensionalized the models’ stress-stretch response by normalizing the stress by the baseline parameter’s matrix modulus value in each model. Further, we set the ratio of *E*_*f*_/*E*_*NH*_ to be 100 for the XIsoComp and OrthoComp models, and set *E*_*f*_/*E*_*HM*_ to be 200 for the XIsoTens model, according to experimental data on sclera [25] and tendon [26].

### 2.5 Multi-start optimization

Parameters were identified by multi-start nonlinear least-squares optimization (NLSQ) with a Monte-Carlo-type approach. We first numerically implemented the five constitutive models (IsoTens, IsoComp, XIsoTens, XIsoComp, OrthComp) in MATLAB, which takes axial deformation and constitutive model parameters as input, imposes the boundary conditions, and returns the Piola stress and deformation gradient tensors as the outputs. Using a fixed set of parameters for each model we calculated the mechanical response and designated this as the baseline response. We then separately performed a multi-start NLSQ optimization routine for each model in an attempt to duplicate this baseline stress response.

To fit the “experimental” axial stress (AS)-axial stretch data, we minimzed an objective function, *f*_*AS*_, defined as

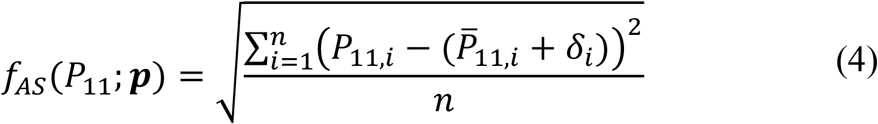

where *P*_11_ is the axial component of the computed stress, 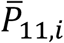 is the corresponding component of the baseline (“experimental”) stress, ***p*** is the set of model parameters, *n* = 20 is the number of discretization steps over the range of *λ*_1_, and *i* is an index identifying the discretization step. In the above, *δ*_*i*_ is a Gaussian noise term (std(*δ*) = 1% of the maximum stress) added to the calculated baseline stress response to mimic experimental noise (e.g., device error). This noise level was chosen based on the expected noise from a commercial load cell, which is commonly 0.25% of the nominal rating of the load cell (see for example [27]) and the assumption that the load cell is operating at 50% of the nominal rating, plus added noise due to other sources (e.g., displacement accuracy and fixture compliance).

The multi-start optimization procedure used the interior-point algorithm (*fmincon*, MATLAB) with a wide search space (Table 1) and 600 random initial guesses (grid size) per parameter. The initial guesses were generated using Latin hypercube sampling, a technique to generate random numbers in a range by avoiding ‘nucleation’ of the numbers (*lhsdesign*, MATLAB).

Any fit with an objective function value of less than 10 times the added noise was accepted as a solution (Criterion 1):

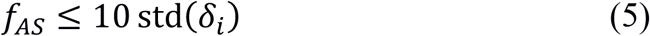

This quantitative criteria was chosen because it provides a consistent basis across the different models for assessing the quality of the fit, and because its upper limit (*f*_*AS*_ = 10 std(*δ*_*i*_)) produces a reasonable fit quality (see supplementary Figure S1).

### 2.6 Including information about lateral strain (LS) predictions

It was immediately evident that the baseline stress-stretch response could be accurately reproduced by many parameter values (see Results), and thus we added a second assessment criterion. We chose this criterion based on the quality of the lateral strain (LS) prediction, because we observed that lateral strain predications were variable across the accepted set of stress-stretch fits. We defined a second objective function, *f*_*LS*_, as

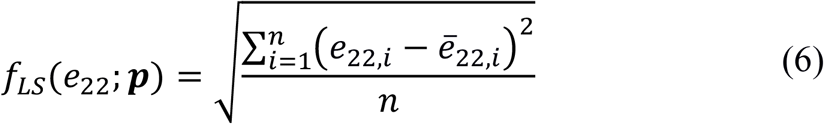

where *e*_22_ is the “22” component of the Lagrangian strain, and other notation is as in equation (4). Similar to the stress fitting criterion, we defined a second criterion for accepting a fit (Criterion 2):

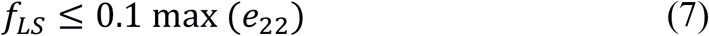

*f* We applied Criterion 2 to the fits that met Criterion 1 (Figure 1), and refer to both criteria together as Criteria 1&2. Note that *f*_*LS*_ was not used to drive parameter optimization; instead it was used to compare the solutions obtained by parameter optimization based on *f*_*AS*_.

Note that the “22” and “33” components of the lateral strain were identical (*e*_22_ = *e*_33_) for most of the models, evaluation of *f*_*LS*_(*e*_22_; ***p***) implies evaluation of *f*_*LS*_(*e*_33_; ***p***). The exception was the OrthoComp model, since the anisotropy of the continuous fiber distribution caused the in-plane deformation to differ along the 2 and 3 axes. Therefore, for this case, we defined a slightly modified objective function for Criterion 2

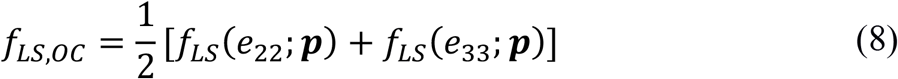

### 2.7 Assessment of identifiability criterion

The identifiability of each material parameter for each constitutive model was assessed by comparing the optimized parameter values to the baseline parameter values. We defined a quantity *γ* to quantify the identifiability of each constitutive model parameter as

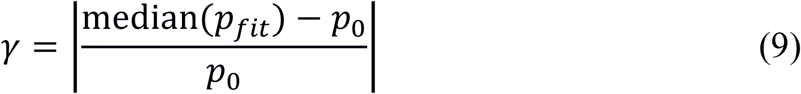

where *p*_*fit*_ is the fitted (optimized) parameter value and *p*_0_ is its corresponding baseline parameter value. A parameter was said to be identifiable if the corresponding *γ* was less than 5%. This threshold was chosen because a 5% deviation is commonly taken as an acceptable error in engineering applications; however, this threshold could be adjusted according to the application (see discussion below).

## 3 Results

### 3.1 Stress fitting results

All the acceptable fitted responses obtained by the optimization of the axial stress (Criterion 1; Eq. 5) closely matched the baseline (“experimental”) stress response (Figure 2). The number of acceptable solutions found through optimization for the IsoComp model was 600/600, i.e. all 600 random initial guesses resulted in fits that satisfied Criterion 1. The corresponding statistics for the IsoTens, XIsoComp, XIsoTens, and OrthoComp models were 252/600, 482/600, 577/600, and 481/600, respectively. However, these fitted responses resulted in non-unique parameter values, i.e. there were many solutions that acceptably reproduced the baseline uniaxial stress-stretch response.

### 3.2 Lateral strain predictions

To investigate the possibility of improving identifiability, we used the lateral strain prediction as a second criterion after optimizing to the axial stress response (Criteria 1&2). Solutions meeting both criteria closely matched both the baseline axial stress-stretch response (Criteria 1&2; Figure 2) and lateral strain response (Criteria 1&2; Figure 3), with multiple acceptable fits overlapping each other and the baseline response (Figures 2 and 3). Importantly, by using both Criteria 1&2, the nonlinear baseline response in XIsoComp (Figure 3C) and the anisotropy in OrthoComp lateral strains (difference between *e*_22_ and *e*_33_) were captured.

**Figure 3:**
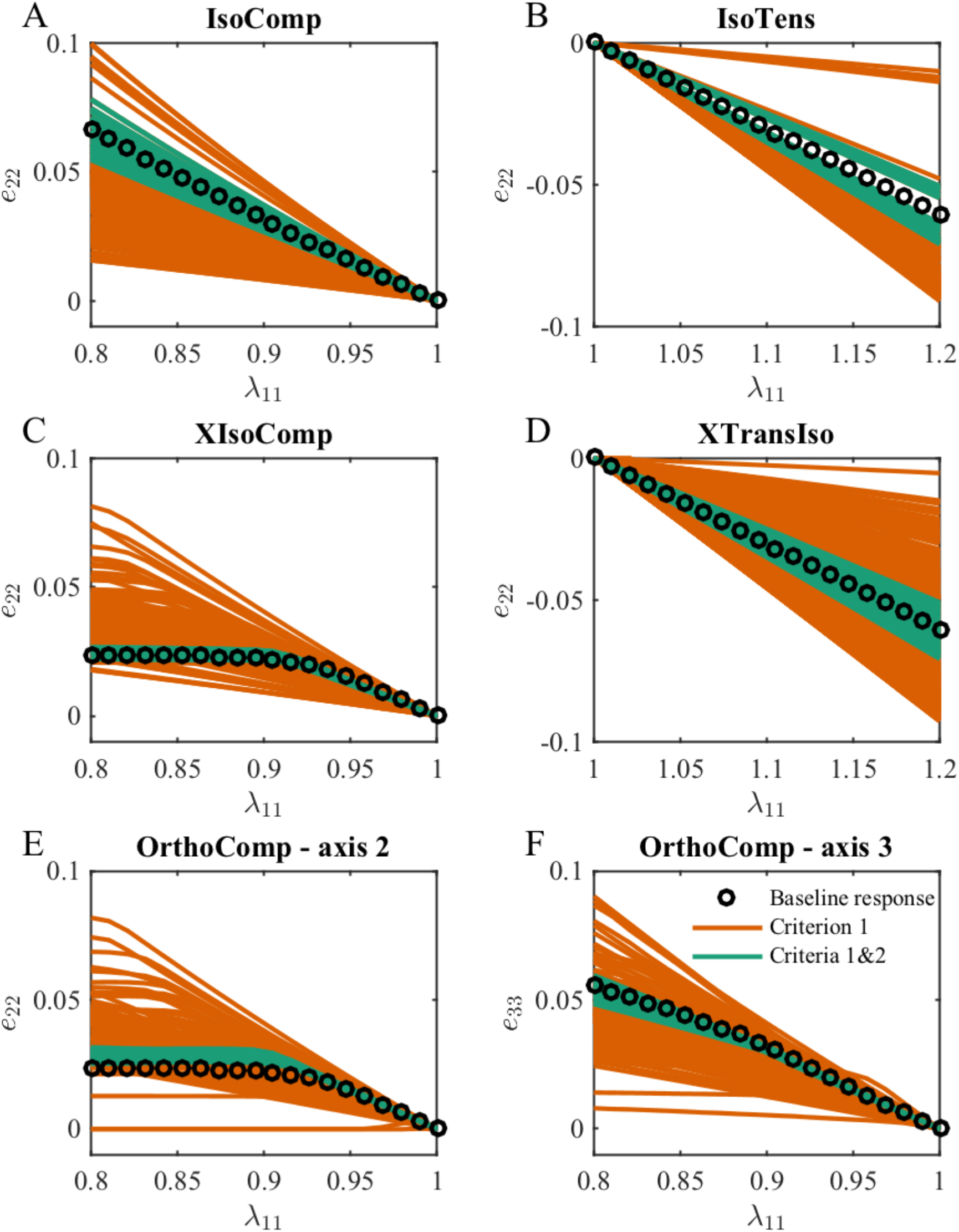
The lateral strain predictions vs. axial stretch based on Criterion 1 (red) and Criteria 1&2 (green) for (A) IsoComp, (B) IsoTens, (C) XIsoComp, (D) XIsoTens, and (E, F) OrthoComp models. We show two plots for the OrthoComp model due to different lateral strain magnitudes in this orthotropic model, i.e. *e*_22_ ≠ *e*_33_. It is evident that using only Criterion 1 yields lateral strain predictions that do not coincide with the baseline (“experimental”) lateral strain, while using Criteria 1&2 significantly improves the match between the fitted results and the baseline values.

The numbers of solutions that met both Criteria 1 and 2 were smaller than the ones identified by only Criterion 1. For example, for the IsoComp model, 600/600 fits were acceptable when using only Criterion 1, while using both Criteria 1&2 reduced this to 56/600. For the IsoTens, XIsoComp, XIsoTens and OrthoComp models, the number of successful fits were 35/600, 19/600, 110/600, and 27/600 solutions, respectively.

### 3.3 Identifiability of parameters

Because multiple solutions were generated with different sets of parameters that matched the axial stress and lateral strain (Figures 2 and 3), we assessed the identifiability of the material parameters by comparing the fitted parameter values to the baseline parameter values. It is convenient to present these comparisons using parallel coordinates plots [16,28], in which each fitted parameter can be compared directly to its baseline parameter value in a single plot (Figure 4). In addition, we quantified identifiability through the quantity *γ* (Table 2), reflecting the material parameter’s error with respect to the baseline parameter value (Eq. 9). In supplementary figures we also provide histograms of the fitted parameter values (Figure S2-4).

**Table 2:**
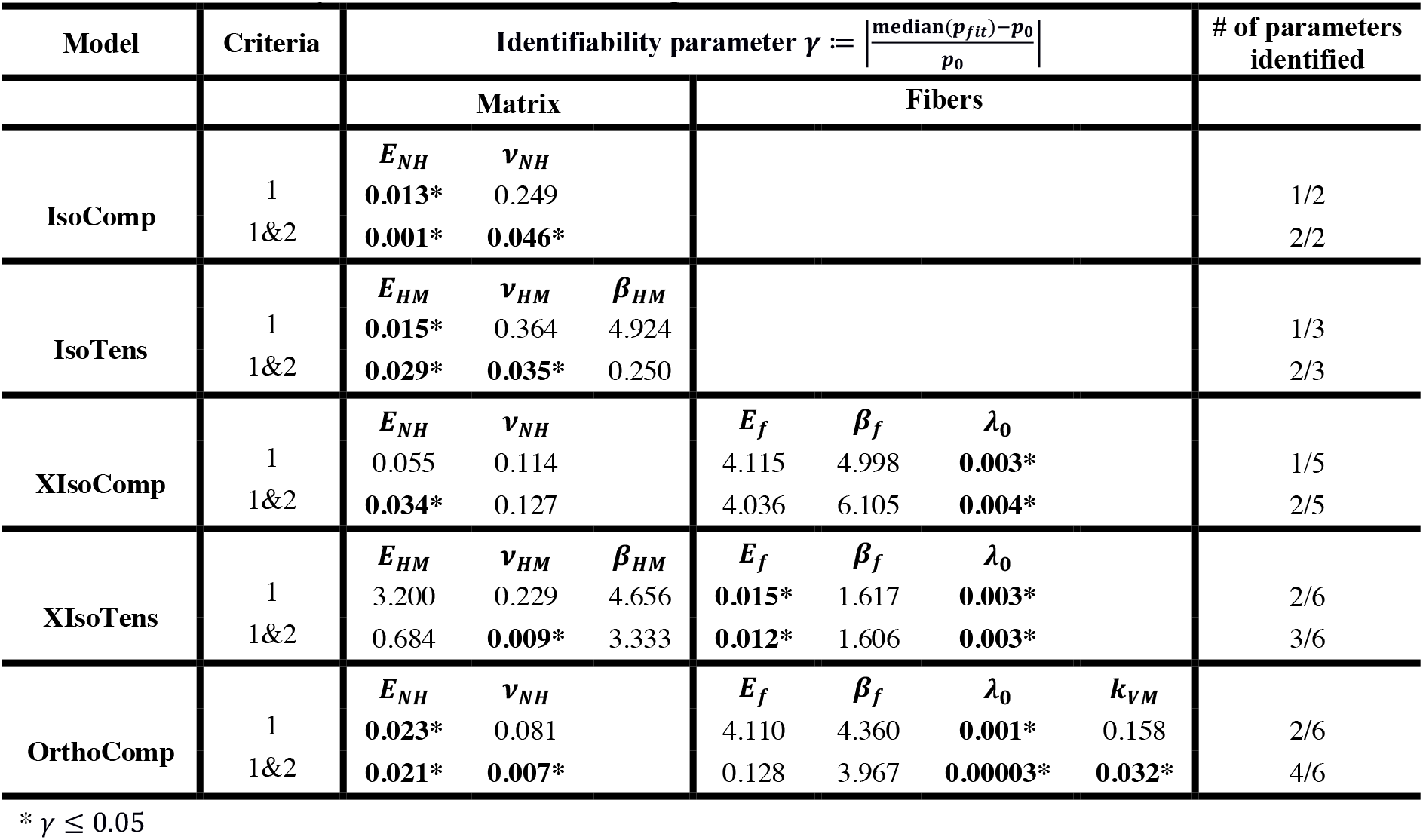
Results of identifiability analysis. A *γ* value ≤ 0.05 indicates that the parameter was successfully identified, where a larger value indicates lack of identification.

**Figure 4:**
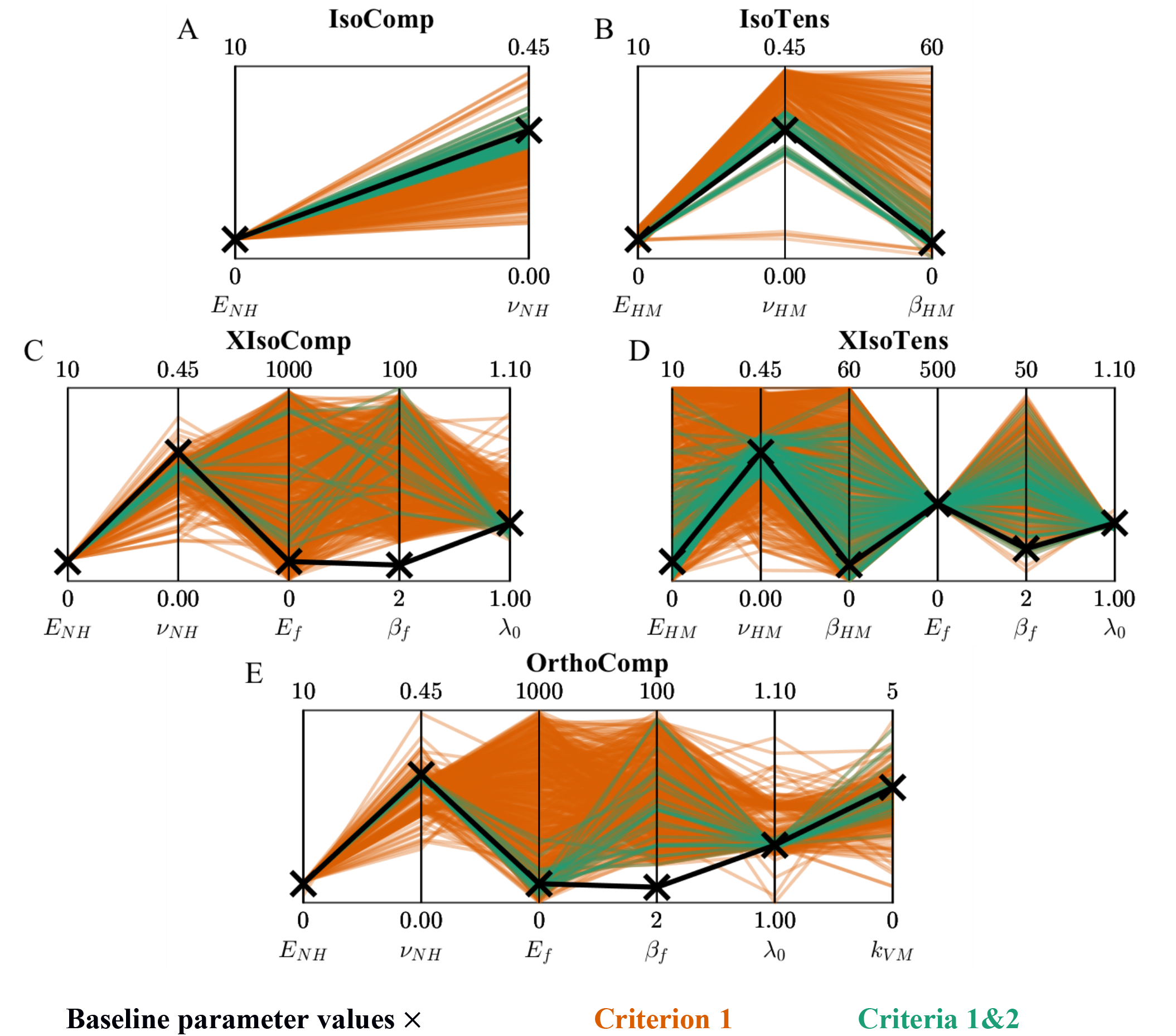
Parallel coordinate representation of the fitted parameter values for the (A) IsoComp, (B) IsoTens, (C) XIsoComp, (D) XIsoTens, and (E, F) OrthoComp models. Fits are shows based on only Criterion 1 (red) and Criteria 1&2 (green). In each plot the vertical lines represent a single model parameter, enabling depiction the high-dimensional model parameters in a single graph.

#### 3.3.1 Isotropic models (IsoComp and IsoTens)

For the IsoComp model, when only considering the stress fits (Criterion 1), the matrix modulus (*E*_*NH*_) was consistently identified (*γ* < 0.05), yet the matrix Poisson’s ratio (*v*_*NH*_) was not (Table 2 and Figure 4A). When considering both stress fits and lateral strain predictions (Criteria 1&2) the deviation from baseline parameter values decreased and both parameters were consistently identified (*γ* < 0.05; Table 2 and Figure 4A).

For the IsoTens model, the modulus *E*_*HM*_ was the only parameter that could be identified when solely matching the stress response (Criterion 1), and this parameter had the smallest deviation from its baseline parameter value among the parameters of the IsoTens model (*γ* < 0.05; Table 2 and Figure 4B). Interestingly, the Poisson’s ratio (*v*_*HM*_) and nonlinearity parameter (*β*_*HM*_) took almost any value in the search range, with the *v*_*HM*_ distribution being skewed toward 0.5 and no meaningful concentration of values for *β*_*HM*_ (Figure 4B). When applying Criteria 1&2, identifiability improved for all parameters; however, *β*_*HM*_ could still not be identified (Table 2 and Figure 4B). The lack of identifiability in the nonlinearity parameter was a common behavior observed in many of the other models (see below).

#### 3.3.2 Transversely isotropic models (XIsoComp and XIsoTens)

As expected, due to there being more parameters in the transversely isotropic models, the distribution of parameter values was larger and more complex than for the isotropic cases. In the XIsoComp model, only the fiber uncrimping stretch *λ*_0_ was identified when Criterion 1 was used (Table 2 and Figure 4C). The matrix modulus, *E*_*NH*_, had a *γ* value 0.055, and despite it having a narrow distribution, it just failed to be classed as identifiable based on our identifiability threshold of *γ* < 0.05. Identifiability was improved by applying both Criteria 1&2, which maintained the identifiability of *λ*_0_ and caused *E*_*NH*_ to become identifiable (Table 2 and Figure 4C). However, values of the fiber modulus (*E*_*f*_) and nonlinearity parameter (*β*_*f*_) were essentially randomly distributed in the search space, and the model was not sensitive to changes in them (Figure 4C).

For the XIsoTens model, none of the matrix parameters were identifiable when considering only Criterion 1 (Table 2 and Figure 4D). However, the fiber parameters, namely fiber modulus (*E*_*f*_) and uncrimping stretch (*λ*_0_), were identifiable and had a narrow distribution (*γ* < 0.05; Table 2 and Figure 4D). Again, the fiber nonlinearity parameter (*β*_*f*_) was not identifiable and had a median value of approximately twice its baseline parameter value (Table 2). When enforcing both Criteria 1&2, the identifiability of the matrix parameters was improved (Figure 4D); however, only *v*_*HM*_ was identified (*γ* < 0.05; Table 2). The fiber parameters remained identifiable and the addition of Criteria 1&2 had almost no effect on their values (Table 2 and Figure 4D).

#### 3.3.3 Orthotropic model (OrthoComp)

When considering only Criterion 1, *E*_*NH*_ was the only matrix parameter that was identified and the uncrimping stretch *λ*_0_ was the only fiber parameter that was identified (Table 2 and Figure 4E). Similar to XIsoComp, the identifiability of fiber parameters was poor, with the fiber modulus *E*_*f*_ and the fiber nonlinearity parameter *β*_*f*_ not identifiable and having nearly uniform distributions over the search range (Figure 4E). The von Mises factor for the fiber distribution (*k*_*VM*_) was centered around 2.5 but did not meet the criterion for identifiability (Table 2 and Figure 4E). Applying Criteria 1&2 improved the identifiability of *E*_*NH*_ and *λ*_0_ as compared to Criterion 1 alone (decreased *γ*), and caused two additional parameters (*v*_*NH*_ and *k*_*VM*_) to become identifiable (Table 2 and Figure 4E). However, fiber parameters *E*_*f*_ and *β*_*f*_ remained unidentifiable, similar to the XIsoComp case.

### 3.4 Identification of Poisson’s ratio

The Poisson’s ratio of the matrix was able to be identified when using Criteria 1&2 for all the models except for XIsoComp (Table 2). Because Poisson’s ratio can be experimentally measured from lateral strain, it was of interest to further investigate identification of Poisson’s ratio. We visualized the relationship between the value of the optimized (fitted) matrix Poisson’s ratio and the values of the objective functions *f*_*AS*_ and *f*_*LS*_, for axial stress and lateral strain, respectively. In general, while *f*_*LS*_ changed by almost two orders of magnitude due to a change in Poisson’s ratio, *f*_*AS*_ was not particularly sensitive to Poisson’s ratio (Figure 5). In compression, *f*_*AS*_ showed some dependence on Poisson’s ratio for the IsoComp, XIsoComp, and the OrthoComp models (Figure 5A, 5C, 5E). However, in tension *f*_*AS*_ was completely insensitive to Poisson’s ratio for the IsoTens and XIsoTens models. In contrast, the curves of *f*_*LS*_ versus *v*_*NH*_ or *v*_*HM*_ showed sharp minima at the respective baseline values (Figure 5A-E) in both compression and tension. We note that the minimum of the *f*_*LS*_ vs. Poisson ratio plot was less pronounced for the XIsoComp and the OrthoComp models (Figure 5C and E).

**Figure 5:**
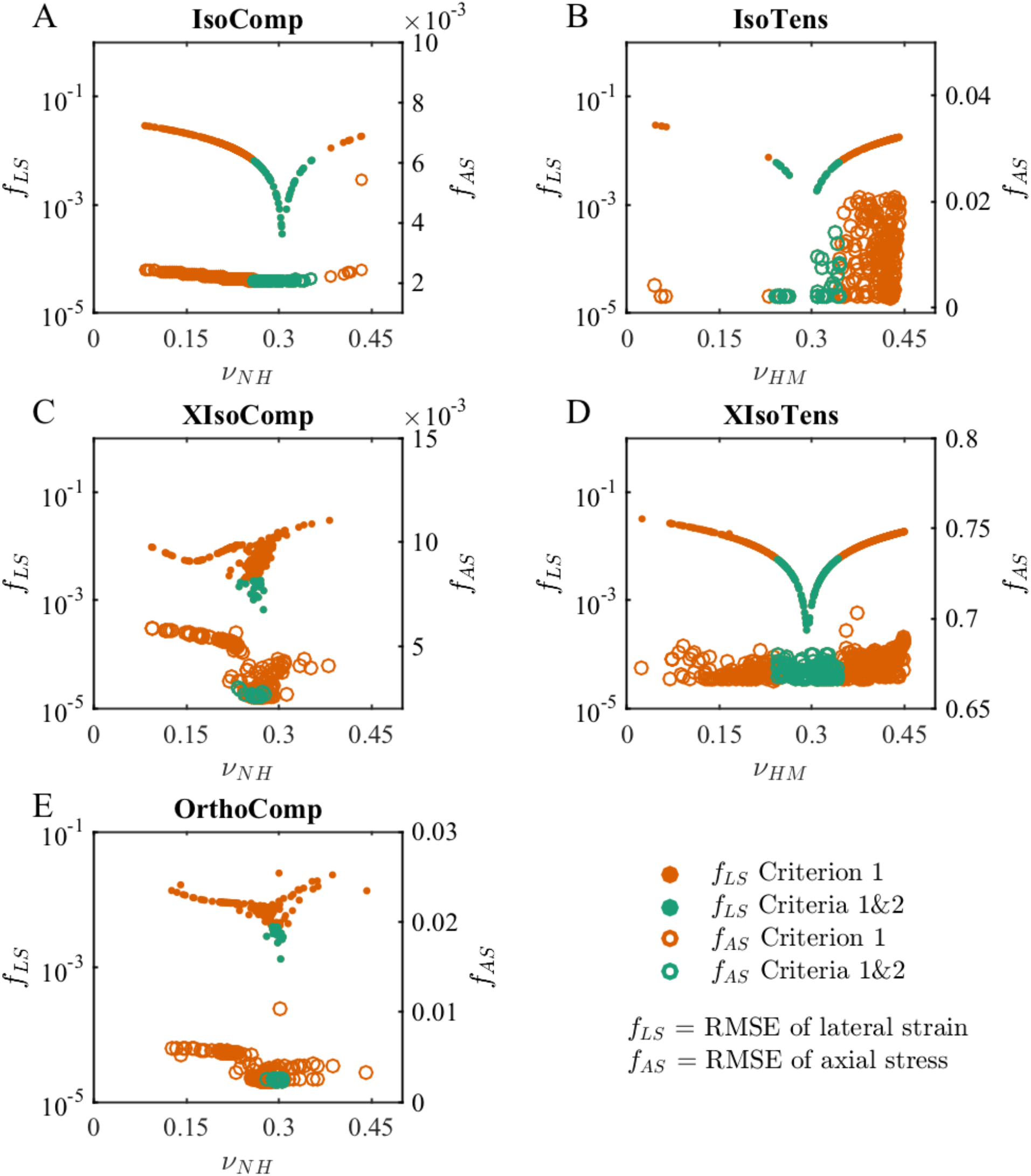
The relation between the objective function values and matrix Poisson’s ratios. We show objective function values for both lateral strain (solid circles; *f*_*LS*_; left axis log-scale) and axial stress (open circles; *f*_*AS*_; right axis linear-scale) for the (A) IsoComp, (B) IsoTens, (C) XIsoComp, (D) XIsoTens, and (E) OrthoComp models. In each model, the fits that met Criteria 1&2 had lower *f*_*LS*_ values compared to fits that only met Criterion 1. However, this trend was not consistent when considering *f*_*AS*_, where only the IsoComp and XIsoComp models (A and C) showed improved fitting with Criteria 1&2. This indicates that although lateral deformation prediction is sensitive to Poisson’s ratio, the same axial stress fits could be achieved with different Poisson’s ratios.

## 4 Discussion

We here investigated the identifiability of constitutive relationship parameters from uniaxial tension and compression mechanical testing of several types of materials: isotropic, transversely isotropic, and orthotropic. These material symmetries and the associated uniaxial testing are relevant to a wide range of applications in fibrous soft tissues, biomaterials, and tissue-engineered constructs. We used baseline material properties relevant for the sclera in compression and for tendon in tension.

Our results showed that, when only the axial stress-stretch response was used in curve-fitting, none of the constitutive models were fully identifiable, i.e. the same stress-stretch response could be generated by different sets of parameter values. We further showed that identifiability of parameters was greatly improved if information about lateral strain was included in the parameter identification.

### 4.1 Identifiability of parameters in isotropic models using uniaxial testing

We considered two isotropic material models; although isotropy is atypical in physiological tissue, isotropic biomaterials, such as hydrogels, are widely used [29]. Our results indicated that even the parameters of the simplest nonlinear constitutive relations were not fully identifiable from a uniaxial test (Figure 4A and B). These results are consistent with studies on the identifiability of parameters in isotropic elastic materials as shown by Hartmann and Gilbert [11]. However, note that the modulus was identified from the stress-stretch response in both isotropic models. Since in many studies, only a measure of stiffness is sought, a uniaxial test could be sufficient for that purpose; however, other material parameters (*v* and *β*) were not identifiable and cannot be relied when deduced by uniaxial testing. Since all these material parameters are needed for finite element analysis, fitting to only stress-stretch data is likely to be insufficient for subsequent use in finite element analysis.

### 4.2 Effect of aligned fiber reinforcement in tensile loading

In the transversely isotropic models in our study, the fiber properties were mostly identifiable, but the matrix parameters could not be identified from only the stress-stretch response. This was particularly noticeable in the transversely isotropic tension (XIsoTens) model (representing tendon, Figure 2D, Figure 4D, and Table 2). Even with addition of lateral strain (Criterion 2) it was not possible to identify matrix modulus (Table 2). This is likely due to the large difference between the stiffness of fibers and matrix in tendon [30]. Other modes of loading such as lateral compression or osmotic loading have been used to remedy this problem [15,26,31]; however, it is likely that in those modes of loading (such as lateral compression) the identifiability of other parameters would be lost (e.g., fiber modulus). A careful *a priori* identifiability check is needed to design the loading mode and analyze the experimental results before relying on optimized material parameters.

### 4.3 Effect of fiber distribution in compression loading

A distribution of fibers perpendicular to the compressive loading direction occurs in several tissues, including the sclera of the eye. When the samples are small, such as in rodent sclera, inverse finite element modeling can be used to determine material parameters, including the spatial distribution of fiber concentration factor, *k*_*VM*_ [4]. Our results showed that, for such anisotropic models in compression (XIsoComp and OrthoComp), the identifiability of parameters is poor (Figure 4 and Table 2), particularly for the fiber modulus and nonlinearity. Surprisingly, the identifiability of the parameters in the orthotropic model (OrthoComp) was superior to that in its transversely isotropic counterpart (XIsoComp). This indicates that anisotropy can improve identifiability of model parameters and highlights the complex interplay between tissue anisotropy and material parameter identifiability.

### 4.4 Improving identifiability of model parameters

It is expected that the addition of more experimental information in data fitting would improve the identifiability of parameters. There are several approaches to include additional experimental data in fitting. For example, experimentally measurement of parameters related to fiber orientation and distribution [24,32] and fiber uncrimping [33,34], and multi-axial mechanical testing [35,36], are potential approaches to improve parameter identifiability. In this study we demonstrated that inclusion of a criterion based on lateral strain when selecting acceptable fits improved parameter identifiability in all the models. However, the most effective approach is not clear and is likely problem-dependent. Examples of other methods include fitting an analytical expression for Poisson’s ratio to experimental lateral deformation [37], or using the direct measurement of Poisson’s ratio, which is appropriate for linear elastic materials and small deformations [11]. However, due to tissue nonlinearity, these methods are not always feasible. The most generalizable options are to either conduct a multi-objective optimization by fitting experimental stress and lateral strain responses simultaneously [38], or to assess the predictions of lateral strain following optimization on the axial stress-stretch response, as was done in this study.

Our analysis of the relationships among matrix Poisson’s ratio and the values of the objective functions for axial stress fit (*f*_*AS*_) and lateral strain (*f*_*LS*_) revealed that, although the stress fit could be practically insensitive to a change in Poisson’s ratio, the lateral strain, as expected, was highly sensitive to Poisson’s ratio and *f*_*LS*_ showed a clear minima near the baseline parameter value for each model (Figure 5). This indicates that, although several values for Poisson’s ratio can result in an acceptable stress-stretch response, only the fits with matrix Poisson’s ratio closest to the baseline parameter value produce a good lateral strain prediction.

### 4.5 Limitations

This study has some limitations. First, although multi-start optimization eliminated dependence of the curve-fitting results on the initial guess, a user-dependent range of parameters is still needed, which could potentially impact the results. However, selecting a wide search space and dense sampling of this space should minimize this effect. Second, we assumed a relatively small noise amplitude in the baseline (“experimental”) stress response. Although this was reasonable for macroscale and tissue testing, in some micro- and meso-scale tests, larger noise levels might be expected, which would affect uncertainty and identifiability. Third, we selected criteria and thresholds for acceptable fits (objective functions *f*_*AS*_ and *f*_*LS*_ based on root-mean squared errors) and for identifiability (*γ* < 5%) which were reasonable for our application. Nonetheless, the threshold and criteria must be selected by the user, and other values could be chosen. Alternative identifiability criteria such as the rank deficiency of the Fisher information matrix are also prone to similar subjective choices of a threshold, where there is a need for definition of a threshold of a “zero eigenvalue” [12]. There are many other examples of identifiability criteria in different disciplines [9,10], which are mostly application driven. Selecting the best identifiability criteria and threshold is problem-dependent and should be adjusted depending on the application.

### 4.6 Summary and conclusions

We investigated the identifiability of material parameters for fiber-reinforced tissues by using nonlinear constitutive modeling and multi-start optimization. Our results indicated that multiple sets of parameters can produce the same stress-stretch responses and that therefore, axial stress response alone is likely to be insufficient to identify tissue material properties. Addition of a selection criterion based on lateral strain significantly improved parameter identifiability; however, even then, some parameters remained unidentifiable. This study is novel in that it provides a systematic approach to assess identifiability of material parameters in a straightforward framework, and the approach is translatable into many studies of parameter identification. Due to the simplicity of implementing multi-start optimization, it could easily be extended to study viscoelasticity and other inelastic behaviors. In conclusion, we recommend conducting an identifiability analysis as a necessary step for any material parameter data-fitting study and suggest multi-start optimization as an effective tool for conducting curve fitting to evaluate the mechanical parameters of tissue.

## 6 Acknowledgments

NIBIB-NIH R01 EB002425, NEI-NIH R01 EY025286, and the Georgia Research Alliance.

## Appendix A: Constitutive relations

### Isotropic models

For the isotropic model in compression (IsoComp) we used a compressible neo-Hookean material description with Helmholtz free energy defined as [21]

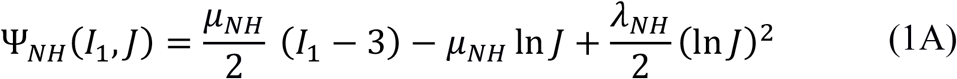

where “NH” stands for neo-Hookean, *I*_1_ is the first invariant of the right Cauchy-Green strain tensor (**C** = **F**^T^ ⋅ **F**; **F** is the deformation gradient tensor) and *J* (the Jacobian of the deformation) is the square root of the third invariant of **C**. The invariants are of **C** are defined in the standard manner as

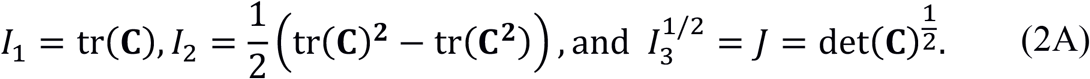

Further, in Eq. 1A 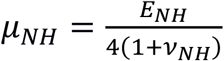 and 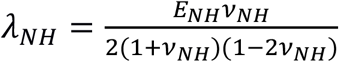 are the Lamé parameters of the matrix, *E*_*NH*_ is the Young’s modulus, and *v*_*NH*_ is Poisson’s ratio. The neo-Hookean constitutive relation has two independent material parameters (*E*_*NH*_ and *v*_*NH*_; Table 1).

For the isotropic model in tension (IsoTens) we used a Holmes-Mow material description with Helmholtz free energy defined as [22]

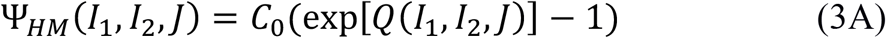

where “HM” stands for Holmes-Mow, and *C*_0_ and *Q* are given by

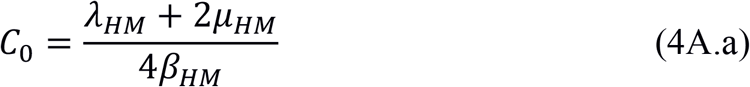

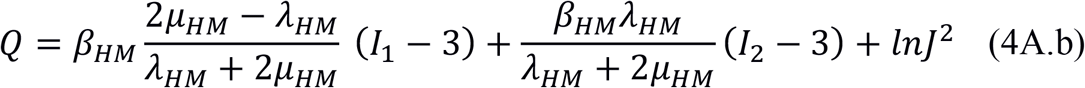

Here, *λ*_*HM*_ and *μ*_*HM*_ are the standard Lamé parameters, *β*_*HM*_ is the nonlinearity parameter (*β*_*HM*_ > 0), and *I*_1_ and *I*_2_ are the first and second invariants of the right Cauchy-Green deformation tensor. For convenience, we show results in terms of Young’s modulus, *E*_*HM*_, and Poisson’s ratio, *v*_*HM*_ instead of the Lamé parameters, which are related to *E*_*HM*_ and *v*_*HM*_ by the following relations: *λ*_*HM*_ = *E*_*HM*_*v*_*HM*_/[(1 + *v*_*HM*_)(1 − 2*v*_*HM*_)], and *μ*_*HM*_ = *E*_*HM*_/[2(1 − *v*_*HM*_)]. This constitutive model has three independent material parameters: *E*_*HM*_, *v*_*HM*_, and *β*_*HM*_ (Table 1).

#### Transversely isotropic models

For the transversely isotropic model in compression (XIsoComp), we embedded a continuous fiber distribution in a compressible neo-Hookean matrix (Eq. 1A) [20]. The fibers obeyed the following formulation

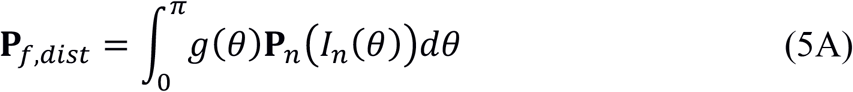

In this relation, **P**_*f,dist*_ is the first Piola-Kirchhoff stress of the in-plane fibers (2-3 plane; Table 1), **P**_*n*_ is the contribution of the fibers oriented with the unit vector ***n*** = cos(*θ*)***e***_**2**_ + sin (*θ*) ***e***_**3**_

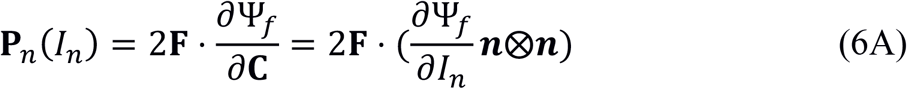

and *g*(*θ*) is the modified von Mises distribution, previously used to describe the fiber distribution in scleral tissue [23,24]

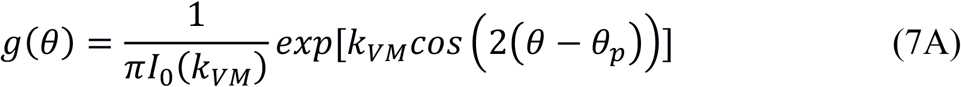

In this relation, *I*_0_ is the modified Bessel function of the first kind, *k*_*VM*_ is the fiber concentration factor (*k*_*VM*_ = 0 for an isotropic in-plane distribution), and *θ*_2_ is the modal fiber orientation. We set *θ*_*p*_ to zero, so that *g*_max_ = *g*(0). In Eq. 6A, Ψ_*f*_ is the Helmholtz free energy of the fiber phase (the subscript ‘*f*’ stands for fibers), defined as:

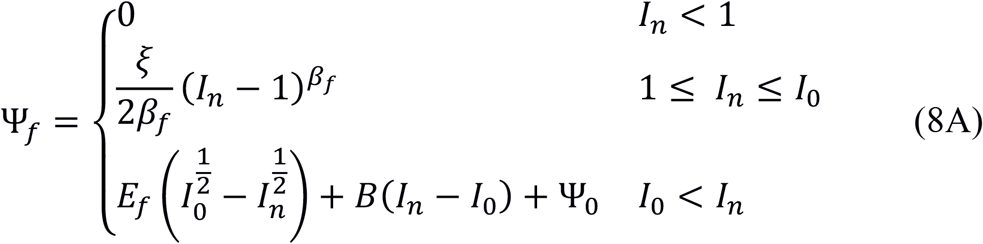

where *I*_n_ is the pseudo-invariant of deformation along a fiber direction defined by unit vector ***n***, i.e.,

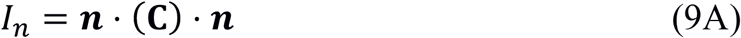

where 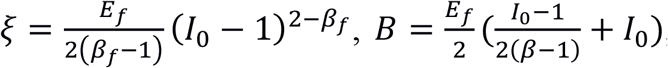, and 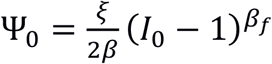

Note that this constitutive relation describes the well-known toe-linear response in tendon [17,18] and this particular form was selected based on its usefulness for 3D finite element simulations [19] (see Section 5.3.7 of the FEBio theory manual v2.9, febio.org). The independent parameters in this constitutive relation for fibers are the tensile fiber modulus, *E*_*f*_, the fiber nonlinearity parameter, *β*_*f*_ (*β*_*f*_ ≥ 2), and the square of the uncrimping stretch of fibers, *I*_0_ = (*λ*_0_)^2^. In summary, there are five independent material parameters in the XIsoComp model (Table 1): [*E*_*NH*_, *v*_*NH*_] for the matrix and [*E*_*f*_, *β*_*f*_, *λ*_0_] for the in-plane fibers, since *k*_*VM*_ is set to zero for the XIsoComp model.

For the transversely isotropic model in tension (XIsoTens), we embedded fibers with a toe-linear response (Eq. 8A) in a Holmes-Mow material (Eq. 3A). The fibers were specified to be parallel to the axial direction (***n*** = ***e***_**1**_). As a result, there are six independent material parameters in the XIsoTens model: [*E*_*HM*_, *v*_*HM*_, *β*_*HM*_] for the matrix and [*E*_*f*_, *β*_*f*_, *λ*_0_] for the fibers (Table 1).

#### Orthotropic model

For the orthotropic model in compression (OrthoComp) we used the same formulation as in (Eq. 5A-9A), except that the fiber distribution was not assumed to be uniform; that is, *k*_*VM*_ > 0. This resulted in an aligned in-plane fiber distribution, and therefore orthotropic symmetry in the material configuration. This senario is a characteristic of prepapillary sclera [23,24]. The OrthoComp model has one more material parameter than the XIsoComp model (six independent parameters): [*E*_*NH*_, *v*_*NH*_] for the matrix and [*E*_*f*_, *β*_*f*_, *λ*_0_, *k*_*VM*_] for the fibers.

---

**Figure S1:**
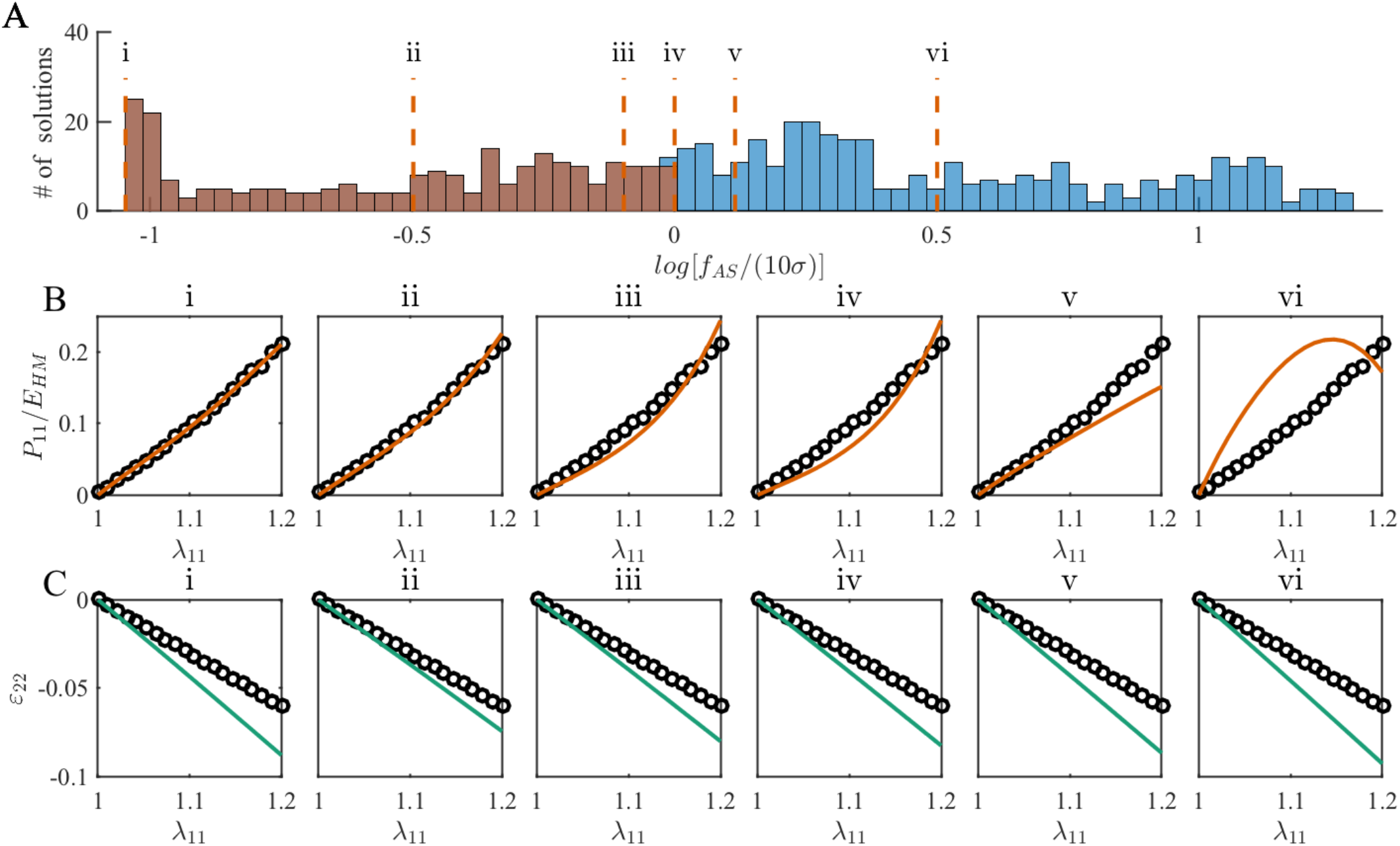
(A) Histogram of values of the objective function *f*_*AS*_ normalized by the threshold value of Criteria 1 for the IsoTens model, where *σ* = std(δ). Note the logarithmic scale on the horizontal axis. Fits with a value less than zero have satisfied Criterion 1 and are shown as brown-colored bars. (B) Examples of selected stress-stretch fits with different *f*_*AS*_ values, as identified in panel A, and (C) their corresponding lateral strain predictions. From the examples it is evident that fits that meet Criterion 1 (i, ii, iii, and iv) have a better fit quality compared to the ones that did not (v and vi). Interestingly, a good quality of stress fit does not indicate a good lateral strain prediction, e.g. compare examples i and ii, where the objective function *f*_*AS*_ was smaller in case i compared to case ii, but the lateral strain predictions were superior in case ii.

**Figure S2:**
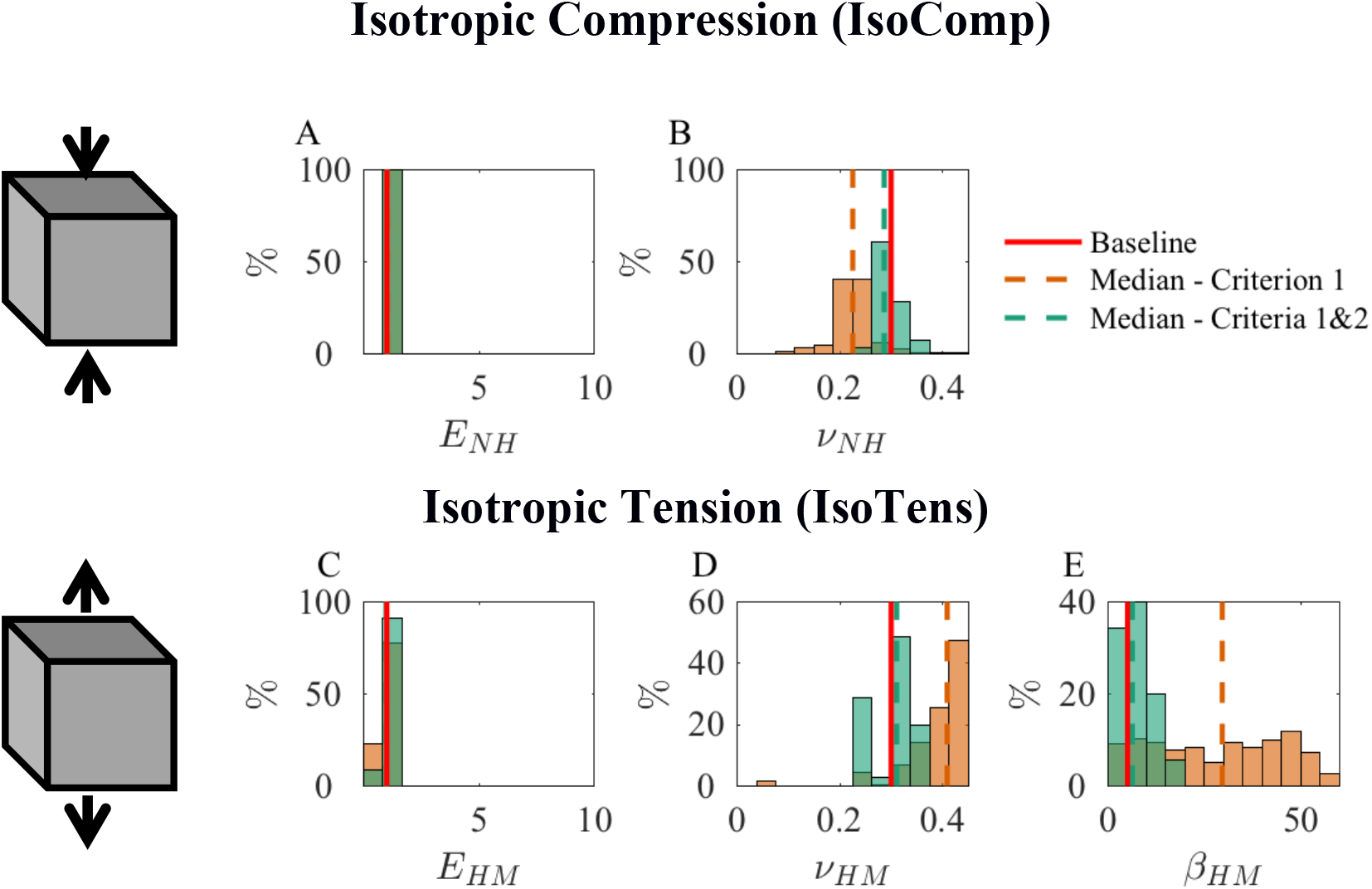
Histograms of fitted parameter values for the (A and B) isotropic compression model (IsoComp), and the (C-E) isotropic tension (IsoTens) model. The vertical red lines show the baseline values for each parameter.

**Figure S3:**
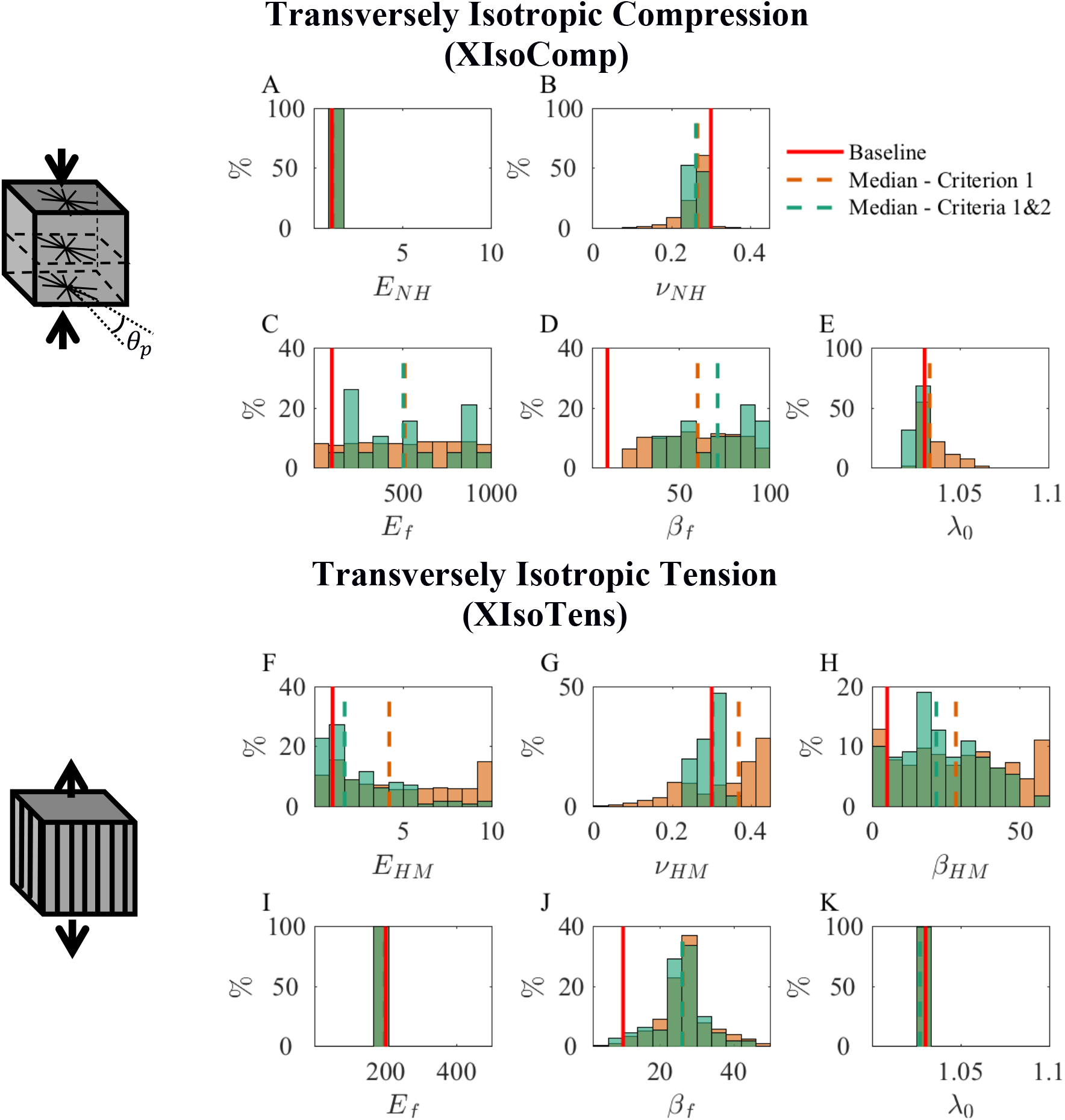
Histograms of the fit parameters for the (A-E) transversely isotropic compression (IsoComp), and the (F-K) the transversely isotropic tension (IsoTens) models.

**Figure S4:**
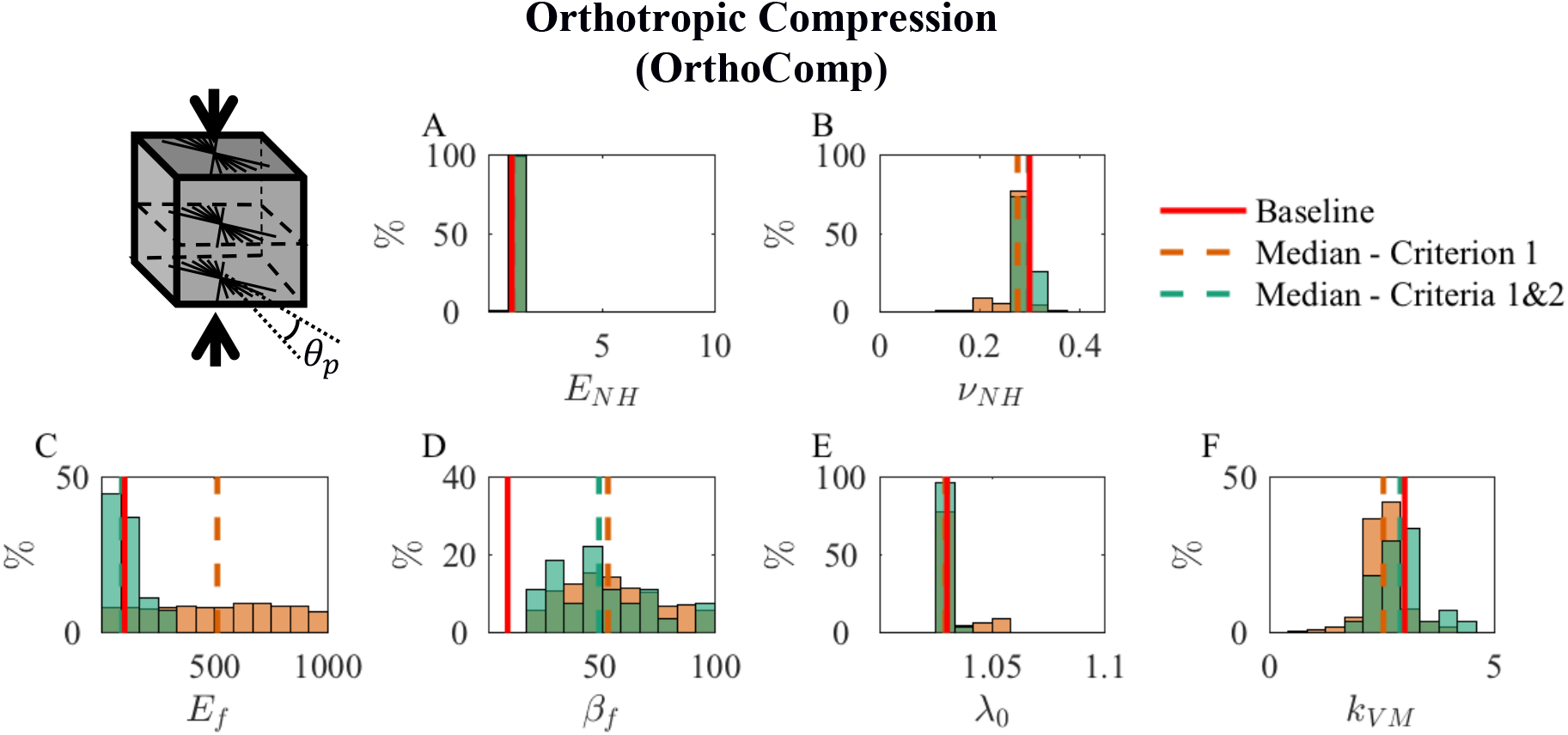
Histograms of the fit parameters for the orthotropic compression (OrthoComp) model.

